# Larger error negativity peak amplitudes for accuracy versus speed instructions may reflect more neuro-cognitive alignment, not more intense error processing

**DOI:** 10.1101/2022.08.19.504504

**Authors:** André Mattes, Elisa Porth, Eva Niessen, Kilian Kummer, Markus Mück, Jutta Stahl

## Abstract

Understanding human error processing is a highly relevant interdisciplinary goal. More than 30 years of research in this field have established the error negativity (Ne) as a fundamental electrophysiological marker of various types of erroneous decisions (e.g. perceptual, economic) and related clinically relevant variations. A common finding is that the Ne is more pronounced when participants are instructed to focus on response accuracy rather than response speed, an observation that has been interpreted as reflecting more thorough error processing. We challenge this wide-spread interpretation by demonstrating that when controlling for the level of non-event-related noise in the participant-average waveform and for single-trial peak latency variability, the significant speed-accuracy difference in the participant-average waveform vanishes. This suggests that the previously reported Ne differences may be mostly attributable to a more precise alignment of neuro-cognitive processes and not (only) to more intense error processing under accuracy instructions, opening up novel perspectives on previous findings.

---

There are hardly any tasks that humans perform flawlessly, i.e. without making any errors. To decrease the number of undesired action outcomes, it is important for the cognitive control system to monitor responses and identify incorrect actions. One marker of response monitoring is the error negativity (Ne; [1], also error-related negativity, ERN; [2]). The Ne is an event-related potential (ERP) component with a fronto-central distribution characterised by a negative peak up about 100 ms after an erroneous response. Although it is usually more pronounced after errors, it can also be observed after correct responses albeit with a smaller amplitude [3]. The Ne is thought to reflect early error processing [1,2] independent from actual error awareness [4]. One of the earliest findings related to the Ne is that its peak amplitude is larger (i.e., more negative) when participants are instructed to respond as accurately as possible compared to when the instruction emphasises speed over accuracy [5]. This finding has been replicated numerous times (e.g. [6–9]). Common explanations suggest that the error significance in the accuracy condition is increased compared to the speed condition, prompting participants to process errors more deeply when accuracy is emphasised [5,10]. A more thorough processing of errors may in turn enhance the participants chances of giving more accurate responses in the future [11]. Here, we present two additional explanations for Ne peak amplitude difference under accuracy and speed instructions that have found little consideration so far: (1) the level of noise and (2) single-trial latency variability.

## Non-event-related noise and averaging

The difference in Ne peak amplitude may be partially attributable to differences in the level of noise included in the averaged waveform. In this context, by noise, we mean random, unsystematic and non-event-related noise which is normally distributed with a mean of zero and a standard deviation reflecting the magnitude of the noise (Figure 1a). This noise is superimposed on the true waveform (i.e. the systematic, event-related signal). The observed waveform thus reflects the sum of the signal and the noise. The principle aim of building a participant-average waveform is to reduce this noise (and thus to increase the signal-to-noise ratio): The more trials go into a participant-average waveform, the less noisy it is. When accuracy is emphasised over speed, participants usually make less errors (e.g. [5,12]). Hence, the participant-average waveform should be based on less trials than when speed is emphasised, and the remaining noise is larger (Figure 1b).

**Figure 1.**
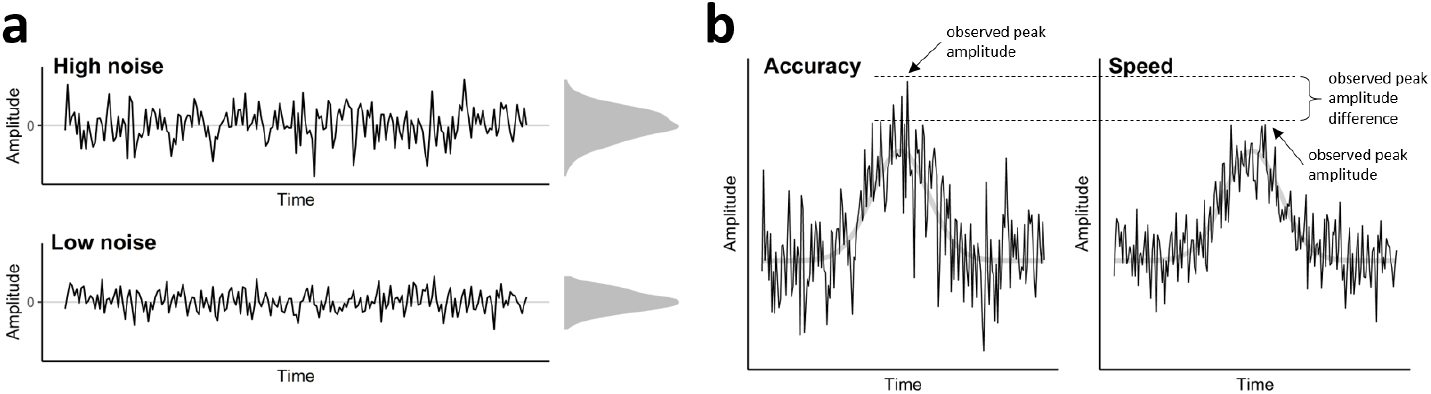
The Impact of Different Noise Levels on the Ne Peak Amplitude. *Note.* A) Different levels of noise after averaging a different number of trials. Both levels have a mean of zero but the standard deviation of the distribution of high noise is larger than the standard deviation of the distribution of low noise. The noise distributions are shown to the right of the plots. B) When the noise is superimposed on the true waveform (light grey), the peak is higher when noise is high than when noise is low.

Often, the peak amplitude is defined as the most positive or most negative point in a certain time interval. This measure not only reflects differences in the true waveform between experimental conditions, but is also highly influenced by different levels of noise (e.g. [13,14]). For example, in Figure 1b, we imposed different levels of random noise on the same true waveform. The peak amplitude in the left panel is much higher than the peak amplitude in the right panel. However, this difference merely reflects different numbers of trials and thus different levels of noise: A lower number of errors in the accuracy condition results in a noisier average waveform and a higher peak amplitude. A higher number of errors in the speed condition results in a less noisy average waveform and thus a lower peak amplitude because the random noise is averaged out.

## Effect of single-trial latency jitter on the peak amplitude

Differences in the peak amplitudes in the participant-average waveforms may also be explained by differences in single-trial latency variability. Peak amplitudes are usually quantified in the participant-average waveform in ERP research because single-trial waveforms are too noisy to reliably detect a peak (e.g. [13]). This approach, however, implicitly assumes that the single-trial waveforms all peak at the same point in time [15]. In Figure 2, we illustrate what happens when this assumption is not met. If the single-trial peak latency variability is large, the average of the single-trial waveforms will have a smaller amplitude than when the latency variability is small. Hence, the observed Ne peak amplitude difference in the participant-average waveforms between the speed and the accuracy condition may also be a result of different levels of single-trial peak latency variability. Specifically, the single-trial waveforms should vary more in their peak latencies in the speed condition than in the accuracy condition, resulting in a flatter participant-average waveform and thus producing a smaller peak amplitude in the average waveform for the speed condition (high latency variability) compared to the accuracy condition (low latency variability).

**Figure 2.**
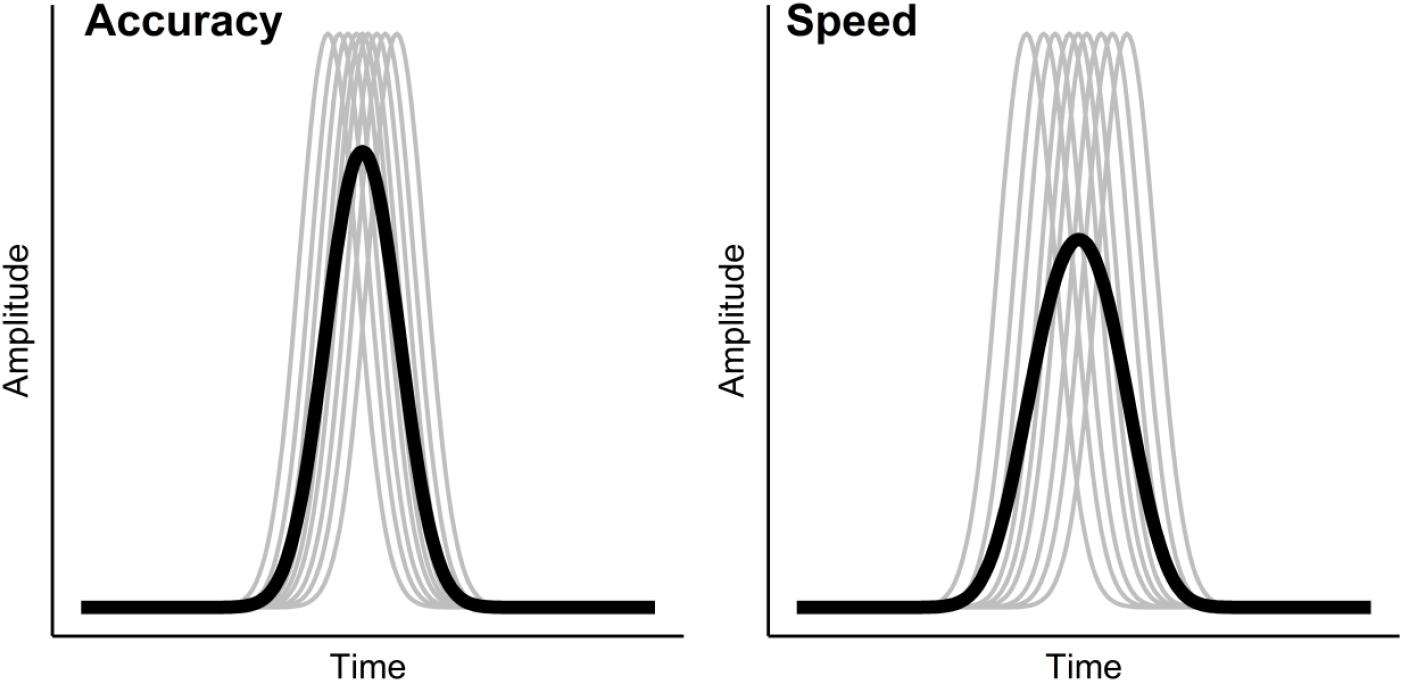
The Impact of Single-Trial Latency Variability on the Participant-Average Waveform. *Note.* The grey lines indicate single-trial waveforms, the black lines result from averaging the single-trial waveforms. In the left panel, the single-trial waveforms peak around the same point in time, hence the variability in single-trial peak latencies is small and the resulting average waveform has a comparatively high peak amplitude. In the right panel, the variability in single-trial latencies is much larger and as a consequence, the resulting average waveform has a lower peak amplitude.

## Objectives

To sum up, we suspect that parts of the difference in the Ne peak amplitude between the accuracy and speed instruction assessed in the participant-average waveform can be explained by difference in the noise level and the single-trial variability in both conditions. Specifically, as we outlined above, we expect that the level of noise (remaining after averaging) is larger in the accuracy condition than in the speed condition, and that the extent of single-trial latency variability is larger in the speed condition than in the accuracy condition. As a consequence, we hypothesize that the Ne amplitude is *negatively* related to the level of noise (the higher the noise, the more negative the Ne peak amplitude) and *positively* related with the extent of single-trial latency variability (the higher the latency variability, the less negative the Ne peak amplitude). (Note that the Ne has a negative peak such that a negative relationship between the Ne and another measure means that higher levels in this measure are accompanied by a more pronounced Ne.) We predict that when controlling for the level of noise and the extent of latency variability, the difference in the Ne peak amplitude between the two instructions should become smaller or even disappear altogether. To test our hypotheses, we analysed existing data of an experiment in which participants received a speed instruction in the first part of the experiment and an accuracy instruction in the second part of the experiment [16–18] and resorted to a single-trial peak estimation technique to assess latency variability.

## Results and Discussion

### Testing Prerequisites

First, we ensured that the instruction had the expected effects in terms of response time and number of error trials. Errors and correct responses were faster in the speed (errors: *M* = 266 ms, *SD* = 104; correct: *M* = 374 ms, *SD* = 94) than in the accuracy condition (errors: *M* = 507 ms, *SD* = 112; correct: *M* = 544 ms, *SD* = 39), *t*(51) = 12.10, *p* < .001, *d_z_* = 2.23, and *t*(51) = 14.20, *p* < .001, *d_z_* = 2.15, respectively, and there was a smaller number of errors when accuracy (*M* = 27.23, *SD* = 19.82) was emphasised over speed (*M* = 86.60, *SD* = 38.17), *t*(51) = 10.59, *p* < .001, *d_z_* = 1.92. (Note that the number of error trials that the EEG analyses were based on was slightly smaller due to the preprocessing of the EEG data, e.g. due to artefact rejection. We report the preprocessing and the resulting trial numbers in the Method section.) Next, we examined our assumption that the averaged Ne peak amplitude was indeed more pronounced in the accuracy *(M* = −13.13 μV, *SD* = 6.91) than in the speed condition (*M* = −8.41 μV, *SD* = 5.28; see also Figure 3a), *t*(51) = 5.16,*p* < .001, *d_z_* = 0.76. As expected, we also found that the level of noise (quantified as the standard deviation of the participant-averaged waveform in the 100 ms pre-stimulus baseline, see Method section) was higher in the accuracy (*M* = 6.27, *SD* = 3.06) than in the speed condition (*M* = 3.56, *SD* = 2.60; see also the density plot at the top of Figure 3b), *t*(51) = 5.02,*p* < .001, *d_z_* = 0.95. Further, the extend of single-trial latency variability was higher in the speed (*M* = 34.42, *SD* = 7.59) than in the accuracy condition (*M* = 27.65, *SD* = 8.33; see also the density plot at the top of Figure 3c), *t*(51) = 5.40,*p* < .001, *d_z_* = 0.85. In conclusion, the instruction had the desired effect in terms of response time and number of error trials, and we replicated the commonly observed pattern of larger Ne peak amplitudes for accuracy vs. speed instructions. Importantly, we were able to demonstrate that the accuracy and speed instruction differed indeed in their level of noise and the extent of latency variability.

**Figure 3.**
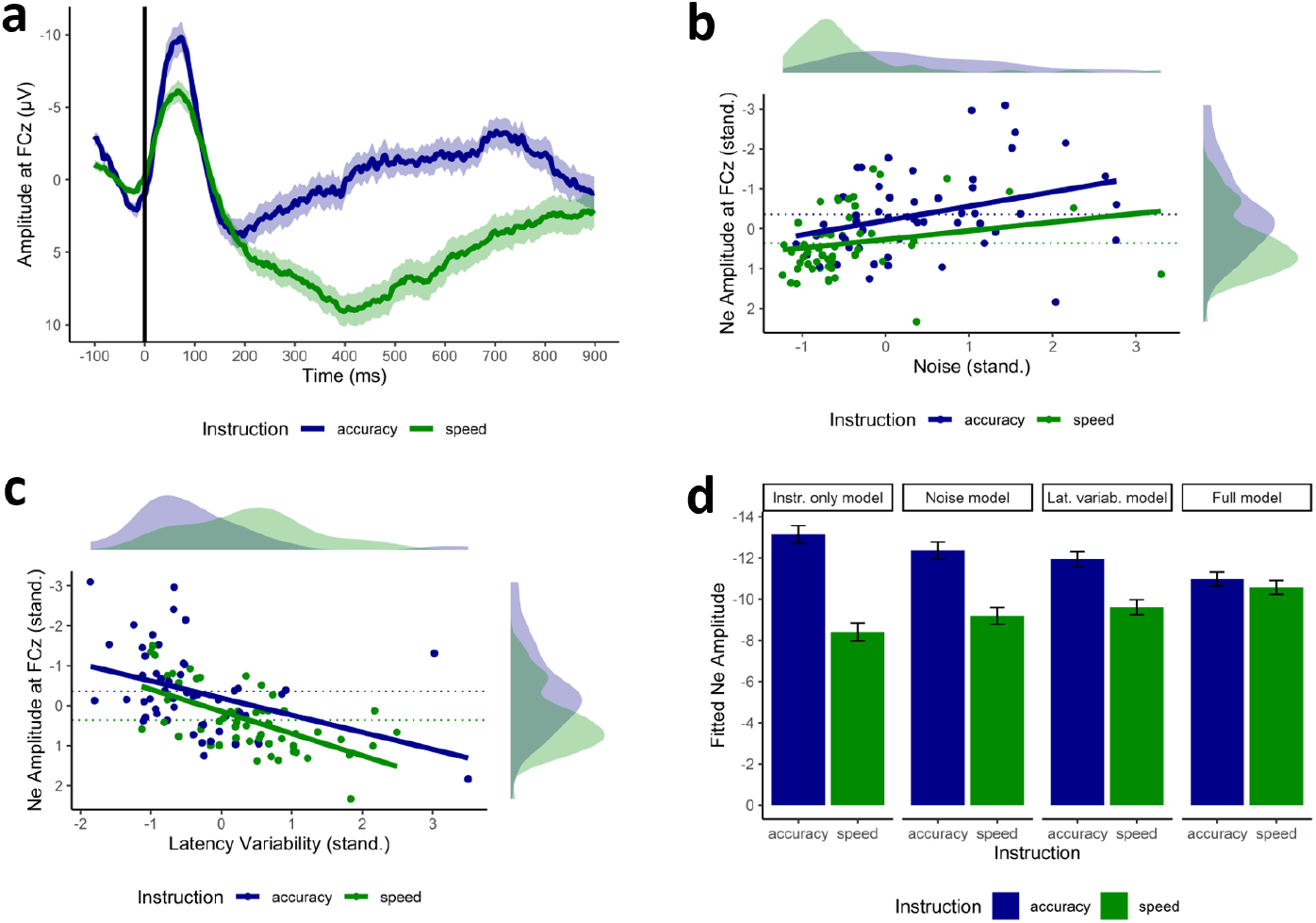
Illustration of the Results. *Note*. A) Grand-average waveform ± standard error of the speed and accuracy condition at the electrode FCz. B) Relationship between the level of standardised noise (x-axis) and the standardised Ne amplitude (y-axis) for the accuracy and speed instruction. C) Relationship between the extent of single-trial latency variability (x-axis) and the Ne amplitude (y-axis) for the accuracy and speed instruction. D) The difference of the Ne amplitude between the accuracy and speed instruction when only considering the instruction (“Instruction only model”), when controlling for the level of noise (“Noise model”), when controlling for the extent of latency variability (“Lat. variab. model”), and when controlling for both noise and latency variability (“Full model”).

### Identifying the sources of speed-accuracy effects on Ne

Next, we tested whether these differences were able to explain the difference in peak amplitude quantified in the participant averages. To this end, we computed a series of mixed models predicting the standardised Ne amplitude. The models are summarised in Table 1. In the first model, we predicted the Ne peak amplitude of the averaged waveform by the instruction only and found that the instruction significantly predicted the Ne peak amplitude, *b* = −0.72, *p* < .001. This analysis is similar to the *t*-test reported in the previous paragraph. Next, we added the level of noise to the mixed model. The level of noise significantly predicted the Ne peak amplitude, *b* = −0.27, *p* = .003. The higher the noise is, the higher (i.e., more negative) the Ne peak amplitude is (Figure 3b). The Ne peak amplitude still differed significantly between the speed and the accuracy instruction, although to a smaller extent, *b* = −0.48, *p* = .002. This finding lends initial support to our assumption that parts of the differences in the Ne amplitude between the instructions can be explained by the level of noise.

**Table 1.** Summary of the mixed models

| Predictors | Model 1 |  |  |  |  | Model 2 |  |  |  |  | Model 3 |  |  |  |  | Model 4 |  |  |  |  |
| --- | --- | --- | --- | --- | --- | --- | --- | --- | --- | --- | --- | --- | --- | --- | --- | --- | --- | --- | --- | --- |
|  | *b* | *SE* | *t* | *df* | *p* | *b* | *SE* | *t* | *df* | *p* | *b* | *SE* | *t* | *df* | *p* | *b* | *SE* | *t* | *df* | *p* |
| (Intercept) | 0.00 | 0.11 | 0.00 | 50 | >.999 | 0.00 | 0.11 | 0.00 | 49 | >.999 | 0.00 | 0.10 | 0.00 | 49 | >.999 | 0.00 | 0.09 | 0.00 | 48 | >.999 |
| Instruction | -0.72 | 0.14 | -5.16 | 50 | <.001 | -0.48 | 0.16 | -3.11 | 49 | .002 | -0.35 | 0.14 | -2.46 | 49 | .015 | -0.06 | 0.15 | -0.41 | 48 | .681 |
| Noise (z) |  |  |  |  |  | -0.27 | 0.09 | -3.03 | 49 | .003 |  |  |  |  |  | -0.31 | 0.08 | -4.00 | 48 | <.001 |
| Latency<br>variability (z) |  |  |  |  |  |  |  |  |  |  | 0.47 | 0.09 | 5.31 | 49 | <.001 | 0.49 | 0.08 | 5.99 | 48 | <.001 |

In the third model, we removed the level of noise as a predictor and entered the extent of single-trial latency variability. As expected, the latency variability positively predicted the Ne amplitude, *b* = 0.47, *p* < .001. The more the single-trial latencies varied, the smaller the Ne amplitude was (Figure 3c). As in the second model, the instruction was still a significant predictor of the Ne amplitude, *b* = −0.35, *p* = .015, although the difference between the instructions was smaller than in the first model, which did not include the latency variability as an additional predictor. This finding provides evidence for our assumption that parts of the differences in the Ne amplitude between the instructions can be explained by the extend of single-trial latency variability.

In the last model, we entered the instruction, the level of noise and the extent of latency variability as predictors. While the level of noise and the extent of latency variability still predicted the Ne amplitude significantly, *b* = −0.31, *p* < .001 and *b* = 0.49, *p* < .001, respectively, the instruction did not reach the level of significance anymore (Figure 3d), *b* = −0.06, *p* = .681. This suggests that the level of noise and the extent of single-trial latency variability were able to explain large parts of the variance in the Ne peak amplitude that is usually found to differ between speed and accuracy instructions. Note however, that the lack of statistical significance of the instruction in the last model does not mean that there is no effect of the instruction on the Ne after controlling for noise and latency variability. The insignificance may also reflect a lack of power to detect an instruction effect which is much smaller than when noise and latency variability are not controlled for *(dz* = 0.06 vs. *d_z_* = 0.72). Importantly, our results show that a higher Ne amplitude for accuracy instructions may not uniquely be interpreted as reflecting a deeper processing of errors due to an increased error significance (e.g. [5,9,10]). It may even be that error processing is just as deep for accuracy instructions as for speed instructions (Figure 3d).

### Level of noise

The level of noise is most likely an artefact due to averaging different number of trials. To avoid artificial differences between the instructions that are uniquely driven by the level of noise, other peak measures which are not as vulnerable to unsystematic noise may be used. For example, when using the mean peak amplitude as the dependent variable in the first model, the effect size is descriptively smaller (peak amplitude: *d_z_* = −0.72; mean peak amplitude (peak ± 20 data points): *d_z_* = −0.51), indicating that the averaging across neighbouring data points reduced the noise in the peak measure (as demonstrated in additional analyses, see Supplementary Table 1). Clayson et al. [14] have discussed the impact of noise on ERP measures extensively, so we will mostly focus on the effects of single-trial latency variability in the following. Unlike the level of noise – which we assume is unsystematic and exclusively due to the different numbers of trials – latency variability potentially provides insights into differences in psychological processes and a different interpretation of the observed speed-accuracy Ne effect.

### Latency variability and process alignment

We find that when accuracy is emphasised over speed, there is less single-trial latency variability in the Ne peak amplitude. This suggests that the cognitive processes following the response are more “aligned” in the accuracy condition. In this context, alignment means the degree to which *across trials*, the same processes occur at the same point in time. The more the processes are aligned, the less single-trial peak latency variability can be observed. In the accuracy condition, time restraints play a less important role than in the speed condition. Hence, at the time of response onset, pre-response processes such as perception, response selection and response initiation are more likely to have finished and subsequent response monitoring processes are executed largely without interferences from previous processes. When speed is emphasised, however, the reduced peak latency variability of the Ne suggests that response monitoring processes may be less aligned across trials, perhaps because pre-responses processes are not fully finished due to the time restraints. Hence, at response onset, different processes might be executed on each trial. Ultimately, these differences in the degree of neuro-cognitive alignment produce more pronounced average Ne waveforms in the accuracy condition (more aligned neuro-cognitive processes) and flatter average waveforms in the speed condition (less aligned neuro-cognitive processes). These interpretations are supported by work by Damaso et al. [19] who differentiate two error types that are linked to accuracy vs. speed instructions. The authors conceptualise the error types in terms of the diffusion model [20] that describes decision making as an evidence accumulation process, i.e. a process in which the information in favour of either of two response options is sampled from the stimulus representation. When accuracy is emphasised, errors may predominantly occur on trials on which the quality of evidence for the correct response is poor. On these trials, spending more time on evidence accumulation would not improve response accuracy [19]. Hence, at the time of response onset, response selection processes have most likely finished and subsequent monitoring processes occur in alignment across trials. When speed is emphasised, errors may predominantly occur because random variations in evidence accumulation prompt a premature motor response [20]. On these trials, spending more time on evidence accumulation in the main task may indeed lead to improved response accuracy [19]. It is thus likely that at the time of response onset, processes such as response selection may continue, postponing response monitoring and introducing latency variability to response monitoring ERPs.

Our analyses provide a different way of interpreting literature findings that build on the instruction effect. For example, Riesel et al. [6] found that patients with obsessive compulsive disorder (OCD) showed a much smaller difference in the Ne amplitude between the speed and accuracy condition than healthy controls, suggesting that patients with OCD have difficulty adapting a response strategy that does not focus on accuracy. Our findings do not change the conclusions Riesel et al. draw from their data, but postulate different underlying mechanisms. Assuming that the same latency effect occurs in their data, we would postulate that while healthy controls seem to disengage from aligning cognitive processes in favour of faster responses when speed is stressed, patients with OCD may have difficulty in doing so. In another study, Endrass et al. [7] found that the Ne amplitude was reduced in older adults compared to younger adults, especially under speed instructions. Hence, for older adults, the speed instruction might disrupt the alignment of cognitive processes even more than for younger adults. These examples demonstrate that speed and accuracy instructions are still useful to study error processing. However, Ne differences may not (only) reflect more thorough error processing or more error significance, but differences in the level of noise and single-trial latency variability. While the level of noise is probably a statistical artefact, the single-trial latency variability may be an alternative mechanism that is helpful in interpreting and understanding differences in the Ne amplitude between speed and accuracy instructions.

### Additional Analyses

We conducted a series of additional analyses to validate our findings and control for potential confounding variables. We analysed the inter-trial phase coherence as an alternative measure of the extent of single-trial peak latency variability. Furthermore, we repeated our analyses controlling for the number of trials. Finally, we recomputed all mixed effects models using the mean Ne peak amplitude (peak ± 20 data points). While these additional analyses corroborated the interpretation of our main results described above, they did not produce new insights that would warrant presenting the analyses in the main text. Hence, we refer the readers to the Supplementary Information for full details on the additional analyses.

## Conclusion

Using single-trial ERP analyses, we could show that substantial parts of the well-known difference in Ne amplitude between speed and accuracy instructions can be explained by differences in single-trial latency variability. This finding shifts the interpretation of the Ne amplitude difference away from processing intensity and neural capacity, and towards temporal dynamics and the alignment of cognitive processes. Our method can easily be applied to research questions in other domains in which the origins of ERP amplitude differences are unknown and may pave the way to novel insights into cognitive mechanisms reflected by ERP amplitude differences.

## Method

### Participants

The study was approved by the ethics committee of the German Psychological Society (DGPs) and was conducted according to the Declaration of Helsinki. In the original (unpublished) experiment, the order of the speed and accuracy instruction was counterbalanced. As the order effect was not relevant for the current investigation, we only analysed the data of participants who completed the speed instruction first and the accuracy instruction second, which was the order with the higher number of participants. Of the 61 participants who completed the task in this order, 9 participants yielded less than six error trials in any of the speed or accuracy conditions (for the most part in the accuracy condition) and were excluded from the analyses (see Electrophysiological Data). The remaining 52 participants (28 male, 24 female) had a mean age of 25.40 years (*SD* = 5.54). All participants gave written informed consent prior to the study.

### Procedure and Experimental Task

Participants performed a modified version of the Flanker Task [21] in which participants had to determine whether a target digit (out of the numbers 1 to 8) was odd or even by pressing a key with one of the index fingers. The assignment of the odd/even responses to the left/right index fingers was counterbalanced across participants. At the left and right of the target digit, two identical distractor digits were presented. Depending on the target-distractor combination, 50% of the presented trials were classified as congruent (i.e. target and distractors are all either odd or even, e.g. 626, 171, etc.) and 50% were incongruent (i.e. the target is odd and the distractors are even or vice versa, e.g. 818, 747, etc.). The stimulus comprising the three digits was presented for 67 ms and was followed by a response window of 1133 ms. Subsequently, a feedback was presented for 700 ms, followed by an inter-trial interval of 1500 ms. In the accuracy condition, the feedback informed participants whether the response was correct or erroneous. In the speed condition, the feedback additionally informed participants that the response was too slow when the response time limit was exceeded. The response time limits were set to 850 ms (accuracy condition) and 85 % of the average response time of the practice block (speed condition). Responses that were given after this response time limit but within the response window were recorded as “too slow”. Participants first completed 10 blocks of 40 trials each in which they were instructed to respond as fast as possible. Next, they completed another 10 blocks in which they were instructed to respond as accurately as possible. More information on the task can be found in Bode & Stahl [16].

### Electrophysiological Data

The EEG was recorded from 61 scalp electrode sites (Fp1, Fp2, F7, F3, Fz, F4, F8, FC5, FC1, FC2, FC6, T7, C3, C3’, Cz, C4, C4’, T8, FPz, CP5, CP1, CP2, CP6, P7, P3, Pz, P4, P8, FCz, O1, Oz, O2, AF7, AF3, AF4, AF8, F5, F1, F2, F6, FT7, FC3, FC4, FT8, C5, IZ, C6, TP7, CP3,CPz, CP4, TP8, P5, P1, P2, P6, PO7, PO3, POz, PO4, PO8) with active Ag/AgCl electrodes (actiCAP, BrainProducts) which were referenced against the left mastoid. A DC converter (BrainAmp DC, BrainProducts) was used to digitise the continuous EEG signal recorded with a sampling rate of 500 Hz. An online band-pass filter (DC-70 Hz) was applied to the signal. The electrooculogram (EOG) was recorded from two electrodes placed at the outer sides of the eyes (horizontal EOG) and two electrodes placed above and below the left eye (vertical EOG).

The EEG data were preprocessed in the *BrainVision Analyzer* software (BrainProducts). First, a DC detrend was applied to the data. Next, the data were stimulus-locked (−100 ms to 2100 ms) and baseline-corrected. An ocular correction was performed [22] and segments in which the signal exceeded ± 150 μV were removed. The stimulus-locked segments were used for the noise estimation (see next section). For the main analyses, the segments were response-locked (−100 ms to 900 ms) and baseline-corrected. After averaging the wavelines within participants, the Ne was defined as the most negative point at the FCz electrode site in the interval from response onset to 150 ms following the response. Only participants who had at least six trials in each of the conditions were considered to ensure a reliable quantification of the Ne [23]. The ERP analyses are based on an average of 23.71 trials in the accuracy condition (*SD* = 18.35, *min* = 6, *max* = 107) and 67.40 trials in the speed condition (*SD* = 28.03, *min* = 15, *max* = 164).

### Noise Estimation

The remaining noise after averaging a given number of trials was estimated as the standard deviation of the participant-averaged EEG signal at the FCz electrode in the 100 ms pre-stimulus baseline. We chose the pre-stimulus baseline over the pre-response baseline because we cannot rule out that systematic processes such as response preparation or even response monitoring are already ongoing before a response is carried out [16]. Unlike the pre-response baseline, we assume that no systematic response-related processes are present in the pre-stimulus baseline and that variation in this interval reflects random noise. The stimulus-locked epochs were baseline-corrected such that the ERP signal of the pre-stimulus interval in which the noise was determined had a mean of zero.

### Latency Variability Estimation

We used a procedure introduced by Hu et al. [24,25] to quantify peak latencies in single-trial waveforms and computed the standard deviation of single-trial latencies for each participant as an indicator of single-trial latency variability. The single-trial estimation was performed in MATLAB (MathWorks) using the *Letswave 6* toolbox [26].

In the following, we briefly describe the single-trial estimation procedure and refer to Hu et al. [24,25] for a detailed description. The procedure consists of two major steps: (1) applying a wavelet filter to denoise the data and (2) modelling the filtered single-trial waveforms in a multiple linear regression approach. In the first step, a wavelet filter is computed by transforming each single-trial waveform from the time domain to the frequency domain and averaging all time-frequency transforms. The resulting average time-frequency transform indicates in which frequencies and at which time points there is stable activation across trials and serves as the basis for the wavelet filter. All single-trial signals are time-frequency transformed, the wavelet filter is applied to them, and they are transformed back to the time domain. The filtered signal is then modelled in a multiple linear regression approach by three regressors capturing the overall waveform, the latency, and the morphology (i.e., whether the waveform is wide or narrow). Finally, the peak is determined in the fitted single-trial waveform and the peak amplitude and latency can be quantified. In a simulation study, Hu et al. [25] have demonstrated that the peak estimates derived from this procedure are accurate and unbiased. The technique has successfully been applied in a multitude of studies (e.g. [18,27,28]). The same trials were used for the participant-average waveform, the estimation of single-trial latency variability, and the estimation of noise.

### Statistical Analyses

In all analyses, we only included errors with response times that did not exceed the response time limit. We used within-subject *t*-tests to investigate differences in response time, number of error trials, Ne peak amplitude, level of noise, and latency variability between the speed and the accuracy condition. We then computed a series of mixed models predicting the Ne amplitude quantified in the participant-average waveform. The models differed in the predictors that they included. The instruction was entered as a contrast-coded dichotomous predictor (speed: −0.5; accuracy: 0.5). The level of noise and the extent of latency variability were entered as standardised continuous predictors. Furthermore, the Ne peak amplitude was also standardized across conditions. This allowed to interpret the regression coefficient of the instruction factor in terms of Cohen’s *d* and the regression coefficients of the continuous predictors similar to partial correlations. Participants were included as random effects in the mixed models. We carried out all statistical analyses in R [29]. We fitted the mixed models to the data using the *lme4* package [30] and tested the regression coefficients for significance using the *lmerTest* package [31]. Data processing and plotting were done using functions provided by the *tidyverse* package [32]. The data and analysis script are available here: https://osf.io/c638e/?view_only=465cdad0831f4d72bdd57fb7bcc4d6ac [anonymous link for review; will be made public upon acceptance of the manuscript]

## Supporting information

Additional analyses

## Declarations

## Acknowledgements

We are grateful to Lina Beck for supporting us in the literature research.

## Data availability

The datasets generated and/or analysed during the current study are available in the OSF repository, https://osf.io/c638e/ [anonymous link for review; will be made public upon acceptance of the manuscript]

