## Additional analyses for "Larger error negativity peak amplitudes for accuracy versus speed instructions may reflect more neuro-cognitive alignment, not more intense error processing"

### Supplementary Information

André Mattes, Elisa Porth, Eva Niessen, Kilian Kummer, Markus Mück, Jutta Stahl

Department of Individual Differences and Psychological Assessment, University of Cologne

### Author Note

This research was supported by the German Research Foundation (STA 1035/7-1).

Correspondence concerning this article should be addressed to André Mattes, Department of Individual Differences and Psychological Assessment, University of Cologne, Pohligstraße 1, 50969 Köln, Germany. E-Mail:

### Supplementary Analyses

To corroborate our key findings and to provide further support for our conclusions, we conducted a series of additional analyses. The key results of these analyses are reported in the main text. Here, we present the results in more detail.

#### Inter-Trial Phase Coherence

We computed the Inter-Trial Phase Coherence (ITPC) of the EEG signal at the FCz electrode site as an alternative measure of neuro-cognitive alignment to validate the finding that there was more single-trial latency variability in the speed condition than in the accuracy condition. The ITPC indicates to which extent oscillations are synchronised across trials (Herrmann et al., 2014; Morales & Bowers, 2022). Hence, if neuro-cognitive processes were more aligned in the accuracy condition than in the speed condition as suggested by our analyses of single-trial peak latency variability, ITPC should be higher in the accuracy than in the speed condition.

**Method.** We preprocessed the EEG data as described in the section “Electrophysiological Data” in the main text and extracted segments ranging from 600 ms before the response to 1000 ms after the response. We conducted the analyses in the EEGLAB toolbox (version 2019, Delorme & Makeig, 2004) for MATLAB (MathWorks). The resulting time-frequency matrix ranged from 109 ms before the response to 507 ms after the response (309 time points), and from 3.4 Hz to 30 Hz (267 linear-spaced frequencies). To transform the EEG signal from the time domain to the time-frequency domain, we set the EEGLAB cycle parameters to 3 and 0.8. This setting resulted in a wavelet that was composed of 3 cycles at the lowest frequency (3.4 Hz) and 5.29 cycles at the highest frequency (30 Hz), with a linearly increasing number of cycles with increasing frequencies. Finally, we performed a baseline-correction using the pre-response interval as baseline. We defined the region of interest (ROI) as ranging from 0 ms to 150 ms and from 4 Hz to 8 Hz. The ROI was based on typical time ranges reported for the Ne (e.g. Danielmeier et al., 2009; Riesel et al., 2019; Tieges et al., 2004) and on literature linking the Ne to the theta frequency band (i.e. 4-8 Hz; e.g. Cavanagh & Frank, 2014; Luu et al., 2004; Yeung et al., 2007). Finally, we averaged the values within the ROI for each participant and each condition.

**Results.** Supplementary Figure 1 shows the results of the ITPC analysis. A visual inspection of the Figure suggested that ITPC was higher when accuracy was emphasised over speed, especially in the time window following the response onset and for frequencies up to 8 Hz. Note that the lowest frequency displayed in the Figure is 3.4 Hz, so it is not possible to draw any conclusions about the ITPC of the delta frequency band (1 to 4 Hz). A *t*-test comparing the ITPC in the ROI between the instruction conditions confirmed that the ITPC was significantly higher for the accuracy instruction ($M$ = 0.44, $SD$ = 0.13) than the speed instruction ($M$ = 0.30, $SD$ = 0.12), $t(51) = 6.74$, $p < .001$, $d_z = 1.11$, supporting the validity of the difference we found in the single-trial latency variability.

**Supplementary Figure 1**

*Inter-Trial Phase Coherence for accuracy and speed instructions*

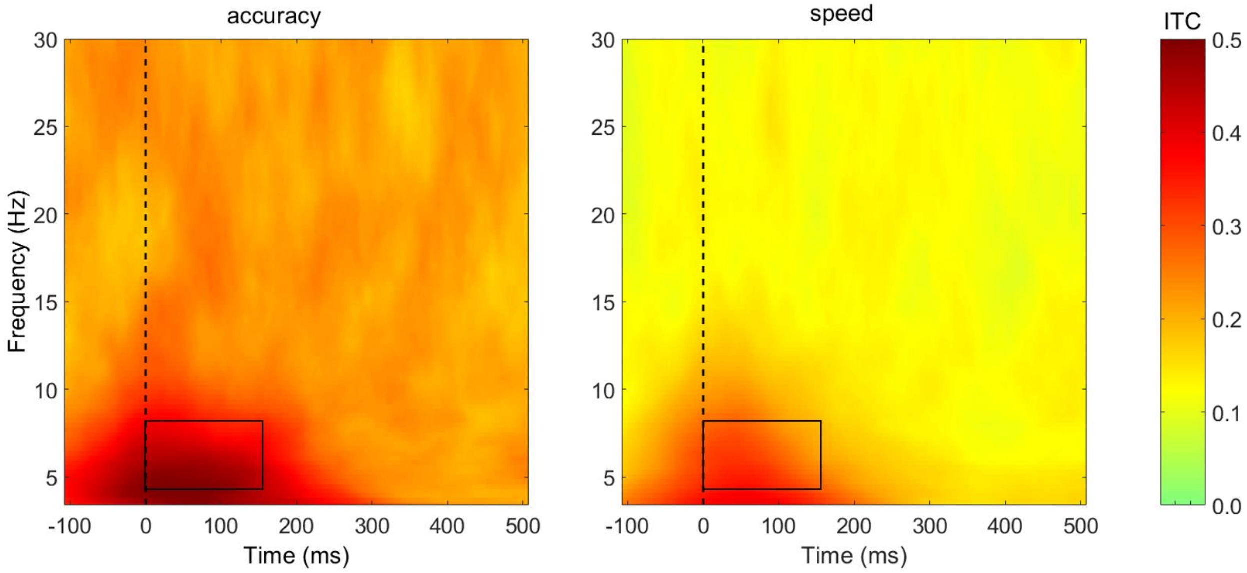


*Note*. The dashed vertical line (time point 0) marks the response onset. The black rectangle indicates the region of interest (ROI) ranging from 0 to 150 ms after response onset and from 4 to 8 Hz (theta frequency band).

**Ne mean peak amplitude**

Considering that the peak amplitude consists of only one data point, it is not surprising that this measure is notoriously distorted by noise. Clayson et al. (2013) have extensively investigated the impact of noise on different amplitude measure and found the mean amplitude to be a more robust measure. We repeated the analyses from the main text for the mean amplitude (peak ± 20 data points). The results are displayed in Supplementary Table 1. Compared to the peak amplitude measure used in the main text, the instruction effect is descriptively smaller (Model 1) and the level of noise is not a significant predictor anymore (Model 2), supporting the claim that using the mean amplitude as a peak measure removes (parts of) the noise. The extent of single-trial latency variability is still a significant predictor and the significant instruction effect vanishes as soon as single-trial latency variability is controlled for (Models 3 and 4).

Interestingly – and unlike in the main analyses – the instruction effect does not emerge in Model 3, either, in which only single-trial latency variability (but not noise) is controlled for. This finding highlights that the impact of noise is (largely) removed by averaging data points around the peak, so noise does not need to be controlled for anymore for the instruction effect to disappear. Overall, these analyses provide further evidence for our claims that Ne peak amplitude differences between accuracy and speed instructions are driven in large parts by (1) different levels of noise and (2) different levels of single-trial latency variability between the instructions.

**Supplementary Table 1**

*Summary of the mixed models prediction the mean Ne amplitude (peak ± 20 data points)*

| Predictors | Model 1 | | | | | Model 2 | | | | | Model 3 | | | | | Model 4 | | | | |
|---|---|---|---|---|---|---|---|---|---|---|---|---|---|---|---|---|---|---|---|---|
| | *b* | *SE* | *t* | *df* | *p* | *b* | *SE* | *t* | *df* | *p* | *b* | *SE* | *t* | *df* | *p* | *b* | *SE* | *t* | *df* | *p* |
| (Intercept) | 0.00 | 0.11 | 0.00 | 50 | >.999 | 0.00 | 0.11 | 0.00 | 49 | >.999 | 0.00 | 0.10 | 0.00 | 49 | >.999 | 0.00 | 0.09 | 0.00 | 48 | >.999 |
| Instruction | -0.51 | 0.14 | -3.60 | 50 | .001 | -0.36 | 0.16 | -2.22 | 49 | .028 | -0.10 | 0.15 | -0.69 | 49 | .494 | 0.10 | 0.16 | 0.61 | 48 | .543 |
| Noise (z) | | | | | | -0.17 | 0.10 | -1.77 | 49 | .080 | | | | | | -0.21 | 0.08 | -2.57 | 48 | .012 |
| Latency variability (z) | | | | | | | | | | | 0.52 | 0.09 | 5.84 | 49 | <.001 | 0.54 | 0.09 | 6.20 | 48 | <.001 |

**Supplementary Table 2**

*Summary of the mixed models prediction the Ne peak amplitude controlling for the number of trials*

| Predictors | Model 1 | | | | | Model 2 | | | | | Model 3 | | | | | Model 4 | | | | |
|---|---|---|---|---|---|---|---|---|---|---|---|---|---|---|---|---|---|---|---|---|
| | *b* | *SE* | *t* | *df* | *p* | *b* | *SE* | *t* | *df* | *p* | *b* | *SE* | *t* | *df* | *p* | *b* | *SE* | *t* | *df* | *p* |
| (Intercept) | 0.00 | 0.10 | 0.00 | 49 | >.999 | 0.00 | 0.10 | 0.00 | 48 | >.999 | 0.00 | 0.09 | 0.00 | 48 | >.999 | 0.00 | 0.09 | 0.00 | 47 | >.999 |
| Instruction | -0.08 | 0.21 | -0.41 | 49 | .681 | 0.01 | 0.21 | 0.04 | 48 | .969 | 0.12 | 0.19 | 0.61 | 48 | .541 | 0.25 | 0.19 | 1.33 | 47 | .186 |
| No. Trials (z) | 0.45 | 0.11 | 4.02 | 49 | <.001 | 0.39 | 0.11 | 3.46 | 48 | .001 | 0.37 | 0.10 | 3.52 | 48 | .001 | 0.28 | 0.10 | 2.74 | 47 | .007 |
| Noise (z) | | | | | | -0.20 | 0.09 | -2.28 | 48 | .024 | | | | | | -0.26 | 0.08 | -3.28 | 47 | .001 |
| Latency variability (z) | | | | | | | | | | | 0.41 | 0.08 | 4.89 | 48 | <.001 | 0.45 | 0.08 | 5.50 | 47 | <.001 |

**Controlling for the number of trials**

There is substantial evidence that the error negativity peak amplitude decreases as the number of errors increases (Wessel, 2018). To ensure that our results were not driven by different numbers of errors in the speed and accuracy condition, we repeated the mixed model analyses and included the number of trials as a covariate, thus controlling for their impact. Note that we used the number of error trials in the behavioural data since these are a better indicator of how many errors a person made than the number of trials in the EEG data. The results of these analyses are displayed in Supplementary Table 2. We found that (similar to findings by Fischer et al., 2017) the number of trials significantly predicted the Ne amplitude in all models. The more errors were made, the smaller was the Ne amplitude. This finding has previously been discussed to reflect differences in the processing of expectancy violation in the anterior cingulate cortex (ACC; Brown & Braver, 2005), the presumed generator of the Ne (Ridderinkhof et al., 2004). In this context, the occurrence of an error represents more of an expectancy violation when the overall number of errors (and thus the error likelihood) is small, triggering more neural activity than when the number of errors is higher and their occurrence is more expected (Fischer et al., 2017; Wessel, 2018). Interestingly, the difference in the Ne amplitude between the accuracy and the speed instruction can be fully explained (statistically) by the different number of errors in both conditions. Even when only the number of trials is included as a predictor in the model, the instruction effect does not reach statistical significance anymore (Model 1). Hence, expectancy violation may play an important role in the context of speed and accuracy instructions. More importantly for our research question, the effects of noise and single-trial latency variability were not affected by the inclusion of the number of trials as an additional predictor in the models. Both of these predictors explained variance in the Ne peak amplitude above and beyond the impact of the number of trials. In fact, the regression coefficient of the number of trials considerably decreased as the other two predictors were included in the model (Model 1: $\beta = 0.45$; Model 4: $\beta = 0.28$), while the regression coefficients of the level of noise and the extent of single-trial latency variability were only slightly smaller than the coefficients in the model that did not control for the number of trials (noise in Model 4: $\beta = -0.31$ not controlling for the number of trials, $\beta = -0.26$ controlling for the number of trials; single-trial latency variability in Model 4: $\beta = 0.49$ not controlling for the number of trials, $\beta = 0.45$ controlling for the number of trials). These additional analyses demonstrate that the effects of the level of noise and – more importantly – the extent of single-trial peak latency variability on the Ne peak amplitude difference between speed and accuracy instructions cannot be explained by different numbers of trials in the speed and accuracy conditions. In fact, the effects of noise and single-trial peak latency variability persist when the number of trials is controlled for.
